# TOC1 supresses PAMP-triggered immunity in *Arabidopsis*

**DOI:** 10.1101/2025.07.16.665052

**Authors:** Olivia J P Fraser, Steven H Spoel, Gerben van Ooijen

## Abstract

The circadian clock synchronizes plants with the rhythmic changes in their environment created by the cycle of day and night. Plant-pathogen interactions are influenced by this rhythmic cycle and a functioning circadian clock is essential for effective resistance to pathogens. In this study we investigate the relationship between PAMP-triggered immune signalling and the circadian clock in *Arabidopsis.* We found that early PAMP-triggered immune responses including flg22- induced *FRK1* transcript levels and the flg22-induced ROS burst are enhanced at subjective dusk rather than subjective dawn. Overexpression of clock gene *TOC1* supressed these defence responses to flg22 and resulted in increased susceptibility to *Pseudomonas syringae*. Additionally, treatment with flg22 and elf18 altered the rhythmicity of *CCA1* and *TOC1.* These results contribute to the currently small amount of research investigating the interactions between the circadian clock and PAMP-triggered immunity. Importantly, we establish the detrimental effect of *TOC1* overexpression on PAMP-triggered defence responses. Further investigation into how clock components regulate early immune responses is essential for improving our understanding of plant health.

## Introduction

Plants, like most eukaryotic organisms on earth, are subject to the roughly 24- hour cycle of day and night. The rhythmic environmental changes caused by this cycle can drastically alter many plant processes. Prediction of environmental changes, due to their rhythmic nature, allows organisms to prepare physiologically or behaviourally to benefit their survival. The circadian clock is an internal timekeeping mechanism that has developed to synchronise organisms with the rhythmic changes in their surroundings caused by the day-night cycle. Environmental cues, such as light or temperature feed into the circadian clock, entraining it to the environment. The clock, in turn, regulates outputs such as gene expression, metabolism, physiology and behavioural processes in roughly 24-hour rhythms ^1–5^.

In *Arabidopsis*, the circadian clock consists of multiple interconnected transcriptional-translational feedback loops, in which circadian components activate or repress each other’s activity in a cyclical manner ^2,4^. The central loop of the clock consists of the dawn expressed MYB-related transcription factors *CIRCADIAN CLOCK ASSOCIATED 1 (CCA1) and LATE ELONGATED HYPOCOTYL (LHY)* and the evening expressed *TIMING OF CAB2 EXPRESSION 1 (TOC1*). *CCA1* and *LHY* act as transcriptional repressors of *TOC1* and of other central clock genes ^6^. *TOC1* also acts as a general transcriptional repressor ^7,8^. Expanding upon the central loop, *PSEUDO-REPONSE REGULATORs (PRRs) 9, 7* and *5* are expressed sequentially throughout the day followed by *TOC1* (also referred to as *PRR1*). All repress *CCA1* and *LHY* and the *PRRs* with earlier phases ^9,10^. Also peaking in expression later in the day, is the Evening Complex (EC) consisting of *EARLY FLOWERING 3 (ELF3), ELF4* and *LUX*. The EC acts as a transcriptional repressor of *PRRs* and *LUX* while being repressed by *TOC1* ^7,11–13^. Positive feedback loops are also present in the clock; *LIGHT-REGULATED WD1* activates *CCA1* and *PRR9* ^14,15^. Additionally, *REVEILLE8 (RVE8)* and *NIGHT LIGHT-INDUCIBLE AND CLOCK-REGULATED1 (LNK1)* and *LNK2* activate expression of *PRR5* and *TOC1* ^16,17^.

The complex rhythmic expression of circadian clock genes regulates many downstream processes in plants, facilitating the temporal separation or coordination of essential processes within the plant in anticipation of environmental conditions ^3,18^. Prediction of environmental changes and potential dangers is essential if plants are to maximise the efficient use of their limited resources. One important system that the clock coordinates is plant innate immunity ^19^.

Plants have evolved a two-layer innate immune system to detect and defend against pathogen attacks ^20^. The first layer of the immune system is triggered upon recognition of pathogen associated molecular patterns (PAMPs) leading to PAMP triggered immunity (PTI). PAMPs are usually highly conserved components found across whole classes of pathogens ^21^. Pathogen recognition receptors at the plasma membrane such as FLS2 detect PAMPs such as flagellin from pathogens ^22^. Upon detecting flagellin, FLS2 forms a receptor complex with BRASSINOSTERIOD INSENSITIVE 1-ASSOCIATED KINASE 1 (BAK1), resulting in the phosphorylation of both FLS2 and BAK1 ^23,24^. This triggers the activation of mitogen-activated protein kinases (MAPKs) which drive the expression of *WRKY* transcription factors and early defence genes such as *FLG22-INDUCED RECEPTOR-LIKE KINASE 1 (FRK1)* within the nucleus ^25^. Alongside this pathway, the plant cell responds to PAMP detection with an influx of calcium ions, stomatal closure, callose deposition and the production of reactive oxygen species (ROS)^26^. Together, these responses lead to resistance to the detected pathogen.

Current research has demonstrated significant circadian regulation of PTI signalling pathway components. In an analysis of publicly available microarray datasets, Bhardwaj et al. (2011) found that pathogen recognition receptor genes *FLS2* and *EFR1* alongside some *MAPK* genes displayed circadian expression patterns under basal conditions. Korneli et al. (2014) found that PAMP-induced expression of *FRK1* was higher at subjective dawn than subjective dusk.

Furthermore, Korneli et al. ^28^ found that PTI-induced production of ROS was higher in the subjective morning versus the subjective evening suggesting circadian control of this process. Basal ROS production and the transcriptional regulation of ROS genes has been shown to be regulated by the circadian clock, with *CCA1* indicated to be key to this relationship ^29^.

Research has also described reciprocal alteration of circadian rhythms by PTI signalling. FLS2 activation shortened the period of *CCA1* rhythms ^30^. Lai et al. (2012) also found that transcription of clock-controlled gene *FKF1* but not *CAB2* was altered by ROS treatments.

In this study, we further examine the interconnections between the circadian clock and PTI signalling in *Arabidopsis* through stimulation of the immune system by bacterial elicitors and virulent bacterial pathogen *Pseudomonas syringae* (*Pst* DC3000). We aimed to investigate the role of core clock genes *CCA1* and *TOC1* in early immune responses, including PAMP-triggered *FRK1* transcript levels and the PAMP-tiggered ROS burst. Additionally, we were interested in how this translated to overall susceptibility to infection with *Pseudomonas.* Our results establish that the clock enhances PTI-induced immune responses at subjective dusk and that *TOC1* overexpression disrupts PAMP-triggered defence pathways.

## Results

### Overexpression of *TOC1* supresses flg22-induced transcript levels of *FRK1* and *FLS2*

Flg22 is a highly conserved 22-amino-acid section of the flagellin protein from *Pseudomonas syringae* that is detected by FLS2 ^31,32^. Elf18 is an 18-amino-acid synthetic peptide derived from the elongation factor Tu protein of *Escherichia coli*, that is detected by pathogen recognition receptor EFR ^33,34^. Following detection of flg22 by pathogen recognition receptor *FLS2*, a cascade of responses results in transcriptional reprogramming for a defence response ^35^. Bhardwaj et al. ^27^ found that *FLS2, EFR,* some MAPK genes and some WRKY transcription factor genes involved in this cascade were rhythmically expressed. To examine the PAMP-induced immune pathway further, we investigated whether the transcript levels of PAMP-induced defence gene *FRK1* exhibited any rhythms. In a single repeat experiment, we measured *FRK1* transcript levels in wild type plants by sampling every 2 hours for two days. Basal *FRK1* transcript levels exhibited visual rhythmicity, with peak transcript levels occurring during the subjective day (Supplementary Figure 1). This suggests potential circadian regulation of basal *FRK1* transcript levels. In a tandem experiment, we treated plants with 1 µM flg22 (Supplementary Figure 1a) or 1 µM elf18 (Supplementary Figure 1b) at 2-hour intervals. We found that flg22-induced *FRK1* transcript levels were also visually rhythmic, however, peak transcript levels occurred during the subjective night. We did not observe rhythmic elf18- induced *FRK1* transcript levels. Rhythmicity analysis of the mean *FRK1* transcript level for each treatment was assessed using the eJTK algorithm ^36^. This analysis revealed that flg22-induced *FKR1* transcript levels are significantly rhythmic (p= 0.02, eJTK_cycle). However, mock and elf18-induced *FRK1* transcript levels are not significantly rhythmic. These results indicate that the circadian clock may gate flg22-induced *FRK1* transcript levels, with responsiveness peaking in the subjective evening and during the subject night. However, more repeats would need to be carried out to confirm rhythmicity or lack thereof in elicitor-induced *FRK1* transcript levels.

To further investigate potential clock regulation of the flg22-induced immune pathway, we compared flg22-induced *FRK1* transcript levels at subjective dawn versus dusk in wild type (*Col-0*), arrhythmic *CCA1ox* plants, arrhythmic *TOC1ox* plants, short period *cca1lhy* mutants and short period *toc1-101* mutants. As expected, wild type plants displayed higher flg22- induced *FRK1* transcript levels at subjective dusk versus dawn (Figure 1, 2). Higher flg22- induced *FRK1* transcript levels at subjective dusk were not consistently observed in either *CCA1ox* or *cca1lhy* plants (Figure 1). This suggests that alteration of clock gene rhythms through *CCA1* affects the temporal difference in flg22-induced *FRK1* transcript levels. Interestingly, *TOC1ox* plants displayed significantly reduced *FRK1* transcript levels at both dawn and dusk after flg22 treatment compared to *Col-0*. *toc1-101* plants did not show consistently higher flg22- induced *FRK1* transcript levels at subjective dusk (Figure 2). Additionally, *toc-101* plants displayed inconsistent flg22-inuded *FRK1* transcript levels compared to *Col-*0, especially at subjective dusk. This disruption could be due to the short period phenotype of *toc1-101* mutants. The low flg22-induced *FRK1* transcript levels in *TOC1ox* suggests that the overexpression of *TOC1* supresses this response of PTI. Further investigation of the flg22- induced defence pathway found that *TOC1ox* plants displayed very little change in transcript levels of *FLS2* after treatment with flg22. *Col-0* and *CCA1ox* plants exhibited significantly increased transcript levels of *FLS2* after flg22 treatment, with *CCA1* displaying significantly higher flg22-induced *FLS2* transcript levels compared to *Col-0* (Figure 3). Together, these results indicate that *TOC1* overexpression supresses multiple elements of the flg22-induced defence pathway.

**Figure 1.**
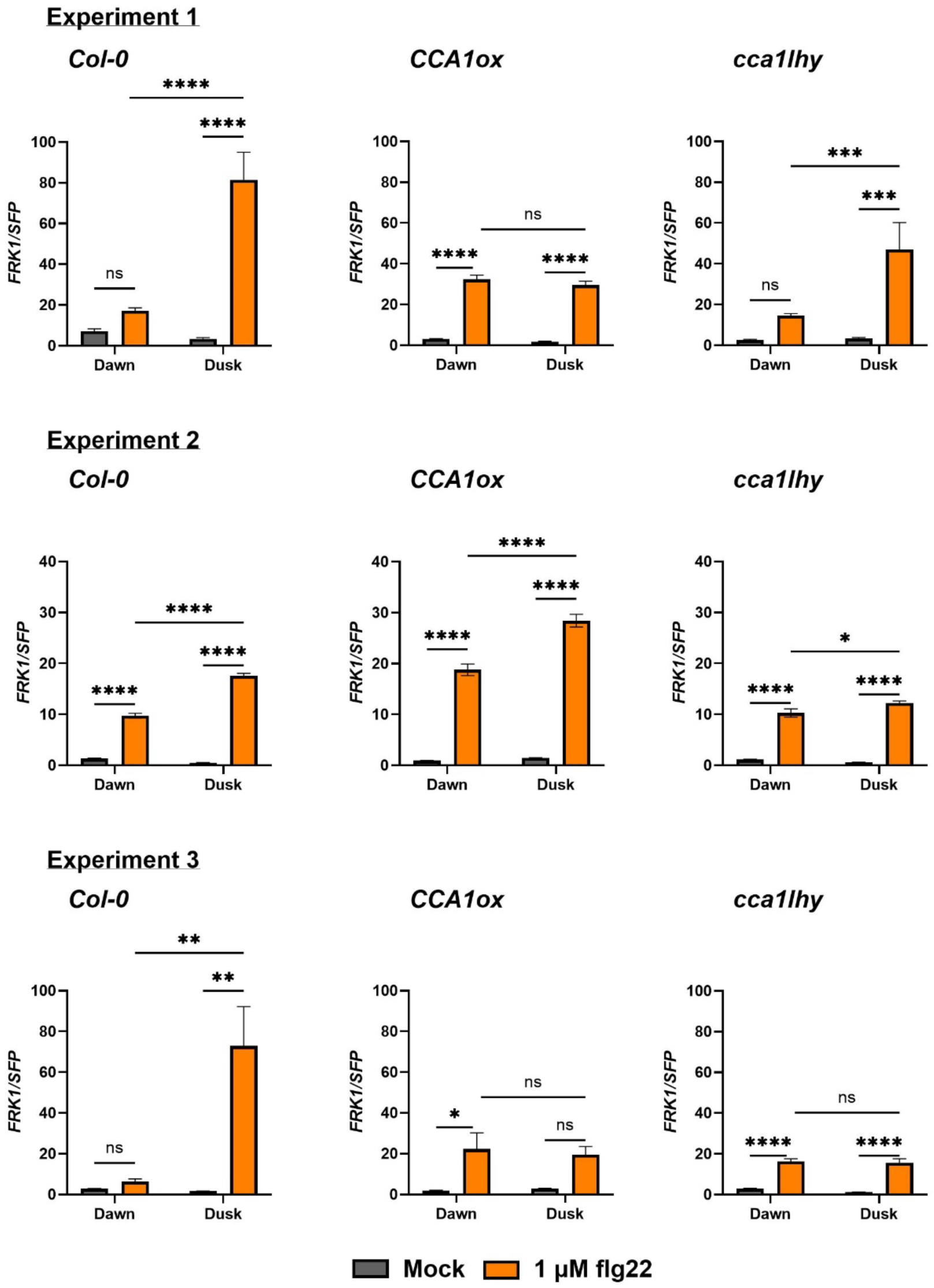
Unregulated expression of *CCA1* disrupts the temporal diAerence in flg22- induced *FRK1* transcript levels. *Col-0, CCA1ox* and *cca1lhy* seedlings were grown on MS plates under split entrainment (12/12 light/dark or dark/light) conditions for 12 days before being released into constant light (LL). At subjective dawn or subjective dusk seedlings were sprayed with 1 μM flg22 or ddH_2_O (mock). Transcript levels of *FRK1* were measured 2 hours post treatment. *p<0.05, **p<0.01, ***p<0.001, ****p<0.0001, two-way ANOVA and Tukey’s HSD. Error bars represent mean ± SEM (n = 4 technical replicates). The data shown here are from 3 independent repeat experiments. All data is shown here to demonstrate the variability between results in *CCA1ox* and *cca1lhy*.

**Figure 2.**
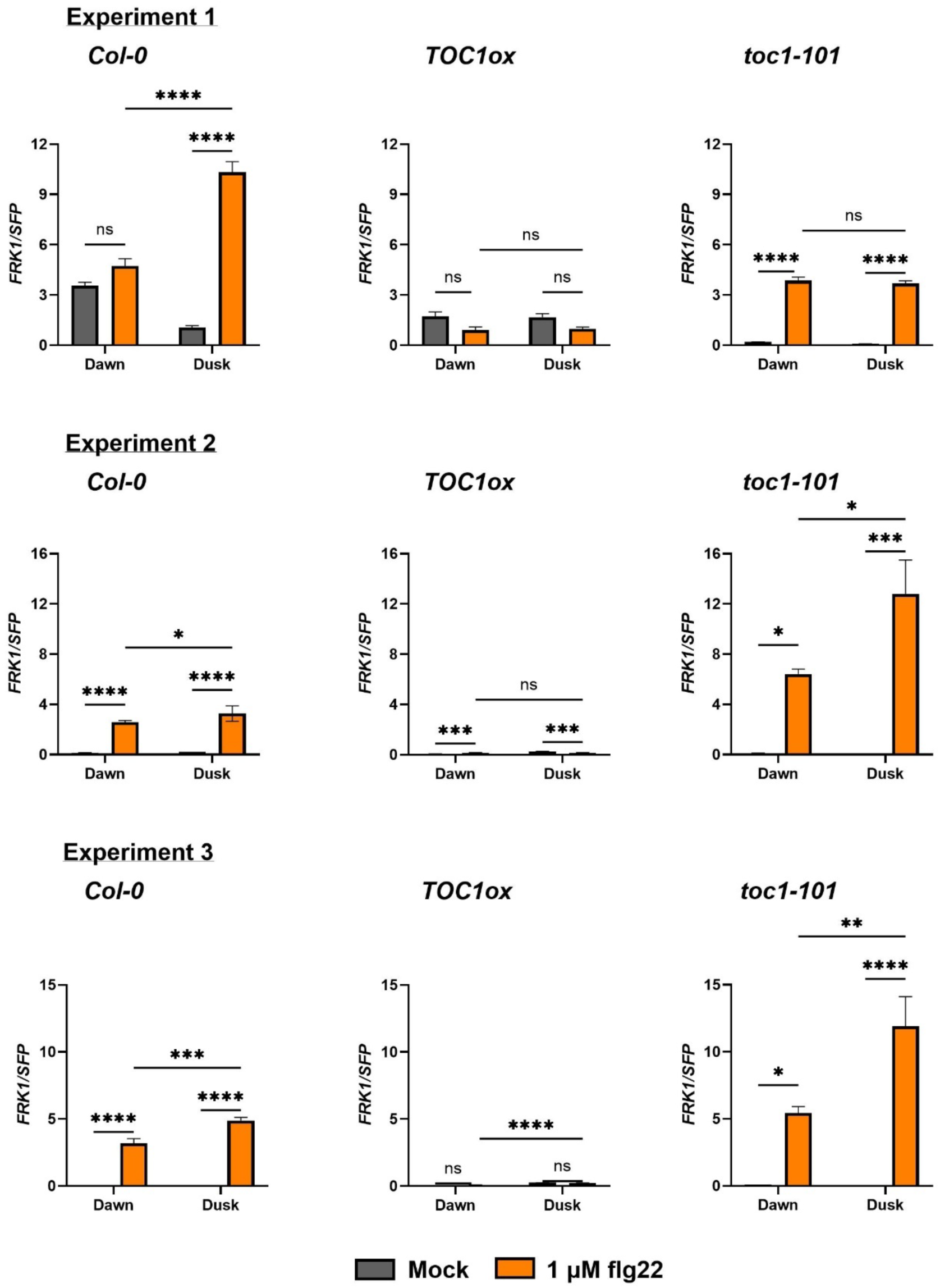
Overexpression of *TOC1* supresses flg22-induced *FRK1* transcript levels. *Col-0*, *TOC1ox* and *toc1-101* seedlings were grown on MS plates under split entrainment (12/12 light/dark or dark/light) for 12 days before being released into constant light (LL). At subjective dawn or subjective dusk seedlings were sprayed with 1 μM flg22 or ddH_2_O (mock). Transcript levels of *FRK1* were measured 2 hours post treatment. *p<0.05, **p<0.01, ***p<0.001, ****p<0.0001, two-way ANOVA and Tukey’s HSD. Error bars represent mean ± SEM (n = 4 technical replicates). The data shown here are from 3 independent repeat experiments. All data is shown here to demonstrate the variability between results.

**Figure 3.**
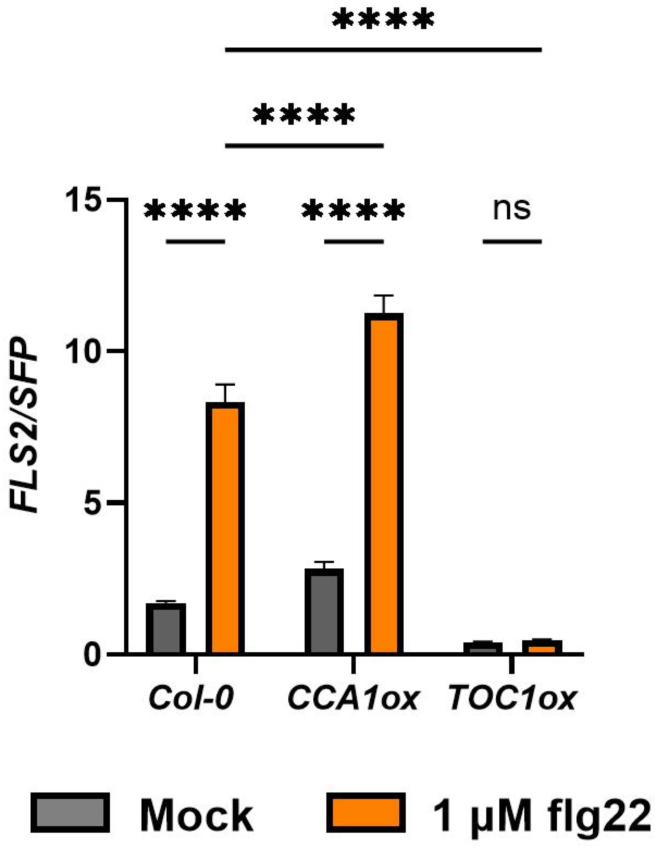
Overexpression of *TOC1* supresses flg22-induced *FLS2* transcript levels. *Col-0*, *CCA1ox* and *TOC1ox* seedlings were grown on MS plates under 12/12 light/dark conditions for 12 days before being released into constant light (LL). At subjective dawn seedlings were sprayed with 1 μM flg22 or ddH_2_O (mock). Transcript levels of *FLS2* were measured 2 hours post treatment. *p<0.05, **p<0.01, ***p<0.001, ****p<0.0001, two-way ANOVA and Tukey’s HSD. Error bars represent mean ± SEM (n = 3 biological replicates). The data shown here are from 3 independent repeat experiments.

### The flg22-induced ROS burst is not observed in *TOC1ox* plants

One of the fastest responses to pathogen detection is the production of the ROS burst. This functions to harm potential pathogens and trigger downstream immune processes in the plant cell ^26^. We measured the ROS burst after treatment with flg22 in *Col-0, CCA1ox, cca1lhy, TOC1ox* and *toc1-101* plants at subjective dawn and dusk. We found that the magnitude of the ROS burst was larger at subjective dusk versus dawn in *Col-0* (Figure 4a, d). This dawn to dusk difference was also observed in *cca1lhy* (Figure 4c, d) and *toc1-101* (Figure 5c, d) mutants, but not in *CCA1ox* plants (Figure 4b, d). This suggests that either clock arrythmia and/or the overexpression of CCA1 affects the circadian gating of the ROS burst. Significantly, we found that flg22 did not induce a detectable ROS burst at either time point in *TOC1ox* plants (Figure 5b, d), indicating that overexpression of *TOC1* supresses the ROS response to flg22 detection.

**Figure 4.**
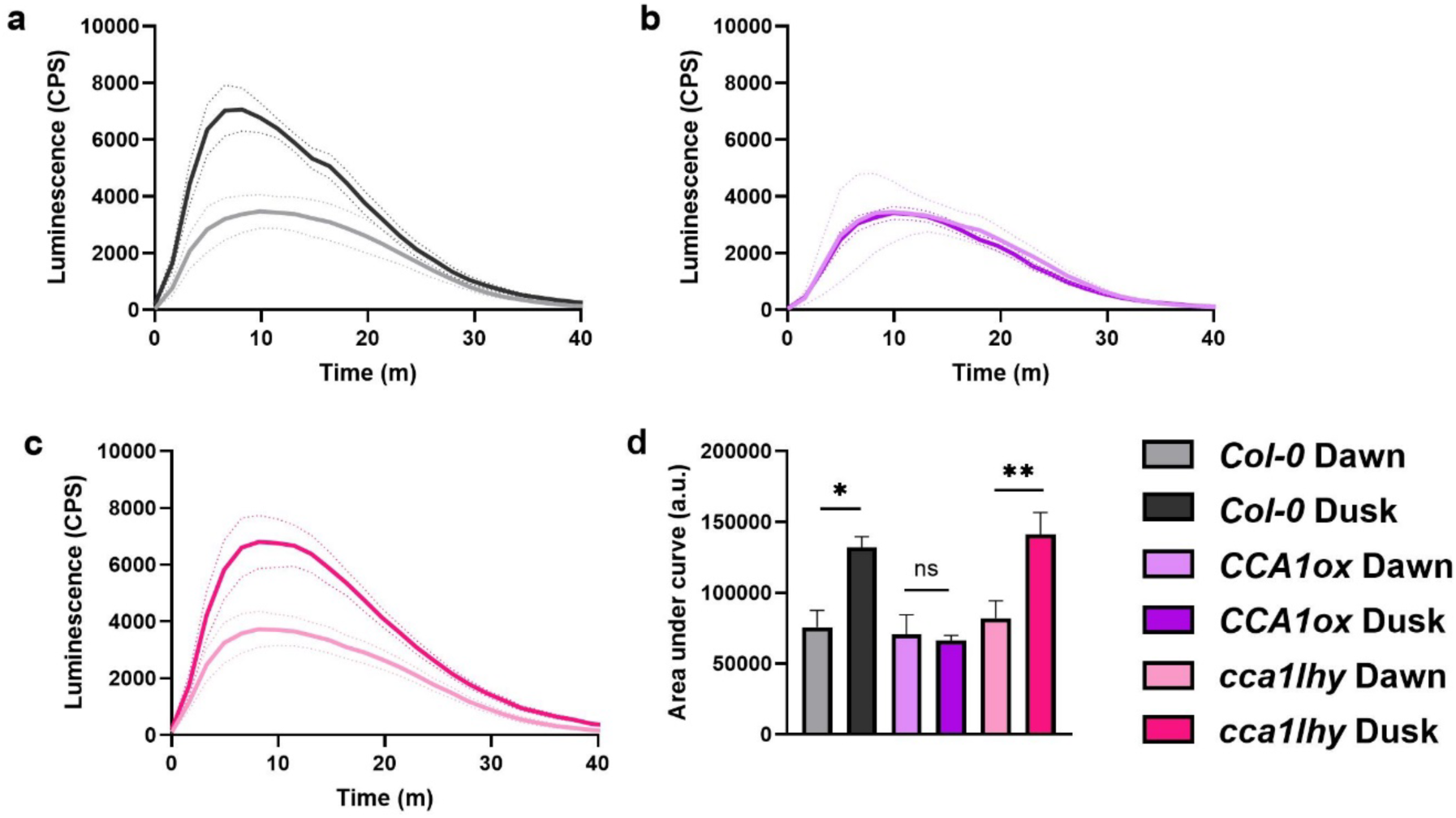
The temporal diAerence in the flg22-induced ROS burst is lost in *CCA1ox*. *Col-0* (a), *CCA1ox* (b) and *cca1lhy* (c) plants were grown on soil under split entrainment (12/12 light/dark or dark/light) for 4 weeks before being released into constant light (LL). After 24 or 36 hours in LL, one leaf disk was cut from each plant and put into the wells of a 96 well plate containing 200 μl of ddH_2_O. 24 hours after leaf disks were cut, the ddH_2_O was removed and an elicitation solution containing flg22 (a-c) or ddH_2_O (not shown) was added. Luminescence was measured immediately. Area under the curve was calculated between 0 - 40 minutes after treatment (d, flg22 treated only). *p<0.05, **p<0.01, ***p<0.001, ****p<0.0001, two-way ANOVA and Tukey’s HSD. Error bars represent mean ± SEM (n = 8). Data shown here are from a single experiment representative of 3 independent repeats.

**Figure 5.**
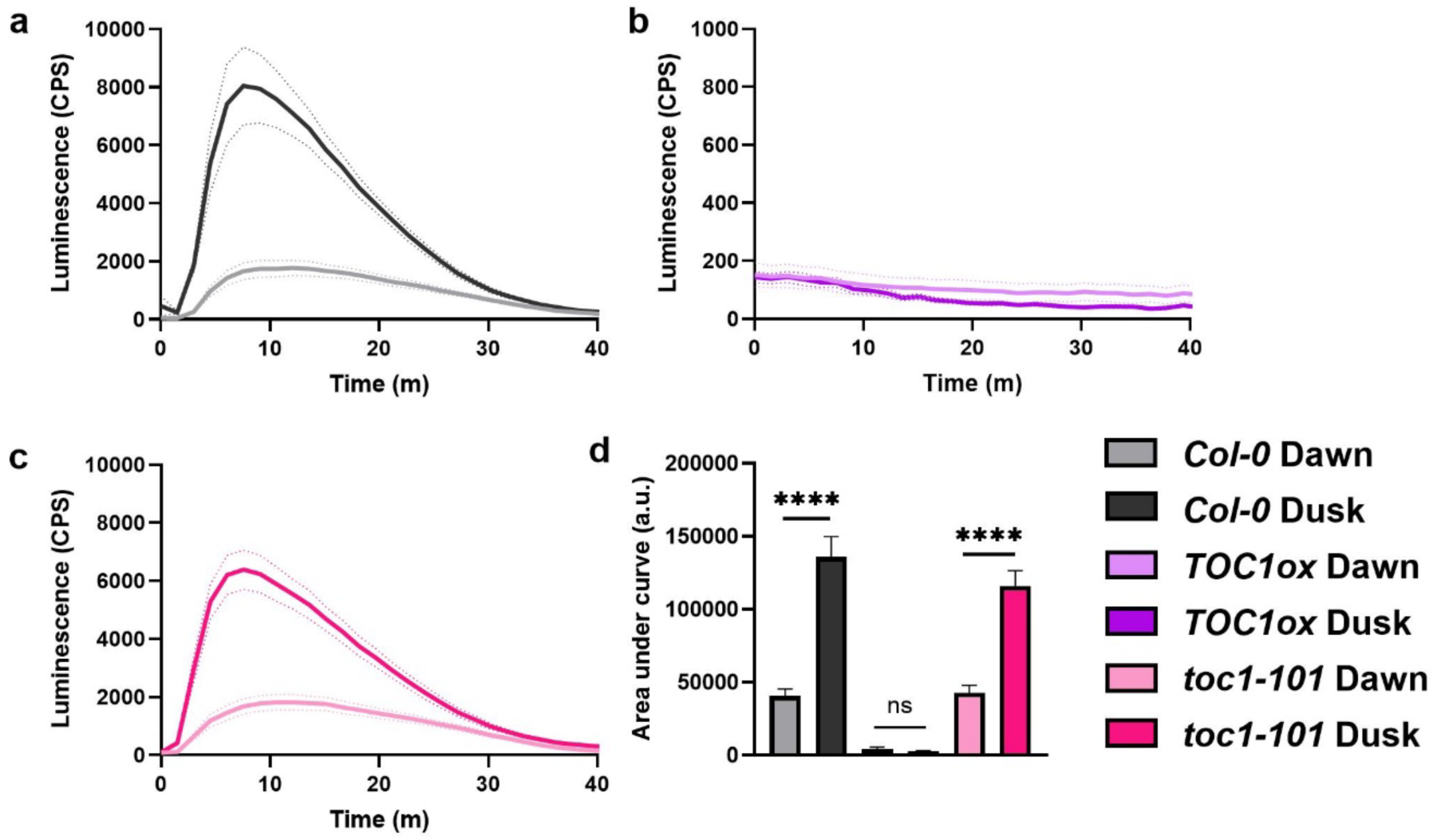
Overexpression of *TOC1* supresses the flg22-induced ROS burst. *Col-0* (a), *TOC1ox* (b) and *toc1-101* (c) plants were grown on soil under split entrainment (12/12 light/dark or dark/light) for 4 weeks before being released into constant light (LL). After 24 or 36 hours in LL, one leaf disk was cut from each plant and put into the wells of a 96 well plate containing 200 μl of ddH_2_O. 24 hours after leaf disks were cut, the ddH_2_O was removed and an elicitation solution containing flg22 (a-c) or ddH_2_O (not shown) was added. Luminescence was measured immediately. Area under the curve was calculated between 0 - 40 minutes after treatment (d, flg22 treated only). *p<0.05, **p<0.01, ***p<0.001, ****p<0.0001, two-way ANOVA and Tukey’s HSD. Error bars represent mean ± SEM (n = 8). Data shown here are from a single experiment representative of 3 independent repeats.

### Overexpression of *TOC1* increases susceptibility to *Pseudomonas* infection

In addition to the ROS burst, PAMP detection leads to an influx of calcium ions, stomatal closure, callose deposition, accumulation of immune hormones and transcriptional reprogramming in an attempt to defend against infection ^26,37–39^. Plant pathogens have evolved to deploy effector proteins which are injected into the plant to modify host function to facilitate a successful infection. Some effectors do this by targeting PTI responses ^40,41^. Detection of these effectors leads to effector triggered immunity (ETI) and the hypersensitive response (HR) in the plant ^42–44^. To investigate the potential influence of the clock on overall PTI, we infected *Col-0, CCA1ox* and *TOC1ox* plants with virulent pathogen *Pseudomonas syringae* pv. *tomato (Pst)* DC3000 wild type and the *hrcC* mutant at subjective dawn and dusk. The *Pst* DC3000 *hrcC* mutant are secretion-defective, thus cannot inject effector proteins into the plant to manipulate infection ^45,46^. The use of the *Pst* DC3000 *hrcC* mutant compared to the *Pst* DC3000 wild type allows the study of PTI with and without the interference of effectors.

Infection with *Pst* DC3000 resulted in increased resistance at subjective dawn compared to dusk in *Col-0.* A similar result was seen in *CCA1ox* plants in some experiments (Figure 6b, d), however, no temporal difference in bacterial growth in *CCA1ox* plants was also observed in other experiments (Figure 6c). *TOC1ox* plants displayed increased susceptibility compared to *Col-0* at subjective dawn, with no significant difference between the subjective dawn and dusk (Figure 6). This indicates that overall immunity to *Pst* DC3000 is increased at subjective dawn in *Col-0* and the overexpression of *TOC1* and *CCA1* disrupt this, though the effect of *CCA1* overexpression appears to be inconsistent.

**Figure 6.**
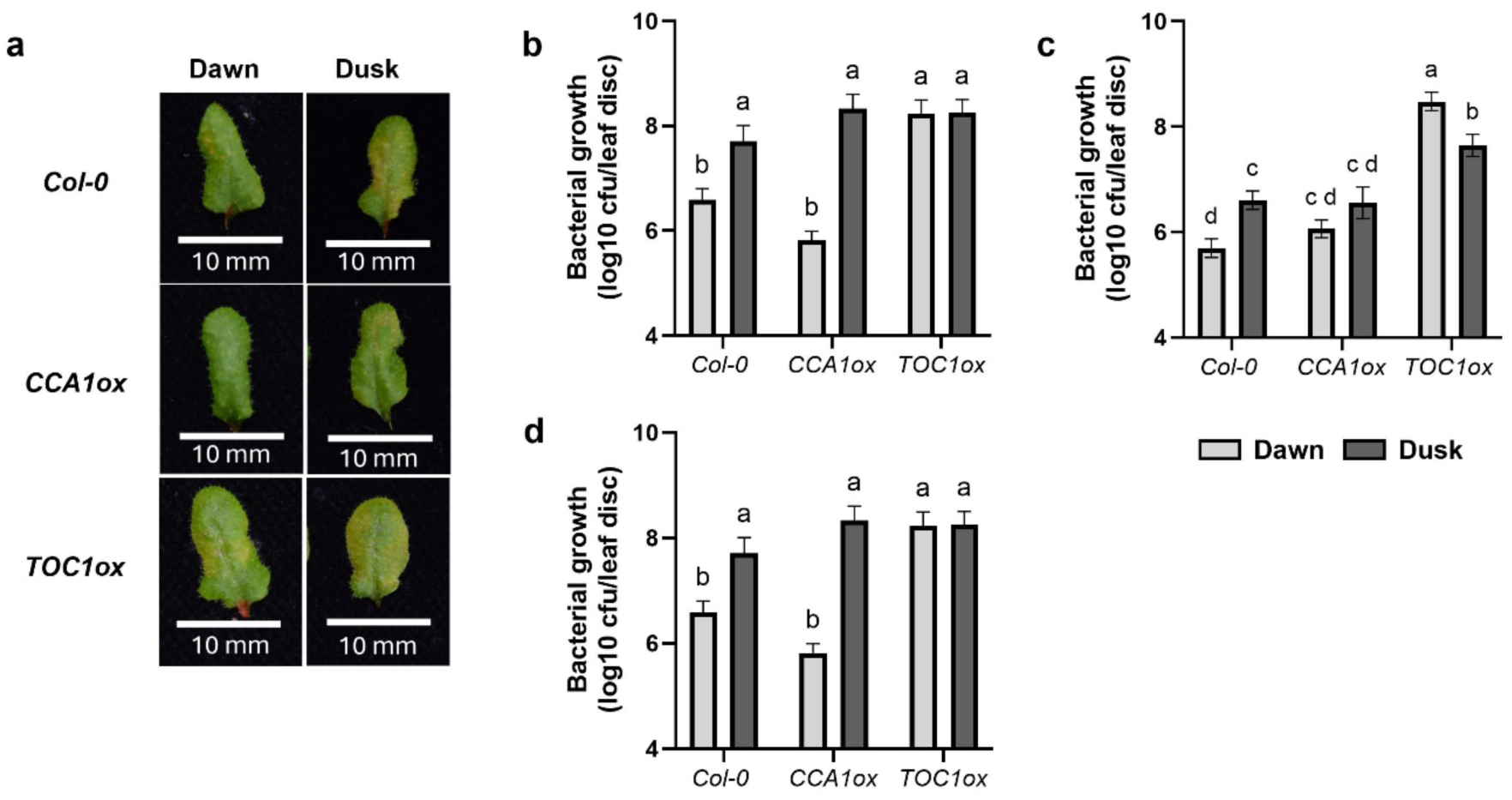
*TOC1ox* is more susceptible to *Pst* DC3000 infection at subjective dawn. *Col-0*, *CCA1ox* and *TOC1ox* plants were grown under 12/12 light/dark conditions for 4 weeks. At dawn on day 28 (LL0) plants were released into constant light conditions. At LL24 or LL36 the plants were infected with *Pseudomonas siringae pv. tomato (Pst)* DC3000. The level of bacterial growth was measured 3 days post infection. (a) Leaves represent the visual median level of infection 3 days post infection for each genotype and time. (b-d) The infection level 3 days post infection expressed by colony forming units (CFU; y-axis is logarithmic) in leaf disk extracts, showing all three repeat experiments in separate graphs. Error bars represent mean ± SEM (n = 8 individual leaves). A two-way ANOVA and Tukey’s HSD Test were preformed to analyse the differences in bacterial growth between time points and genotypes. The letters indicate significant difference between groups (p < 0.05). The data shown in (a) and (b) are from the same experiment. (c) and (d) are data from independent repeat experiments.

*Col-0* infected with *Pst* DC3000 *hrcC* did not display a significant difference in resistance between subjective dawn and dusk. *CCA1ox* plants exhibited similar levels of bacterial infection to *Col-0.* However, *TOC1ox* plants showed increased levels of bacterial growth compared to *Col-0* (Figure 7), indicating that plants overexpressing *TOC1* have impaired PTI during infection with *Pst* DC3000 *hrcC*.

**Figure 7.**
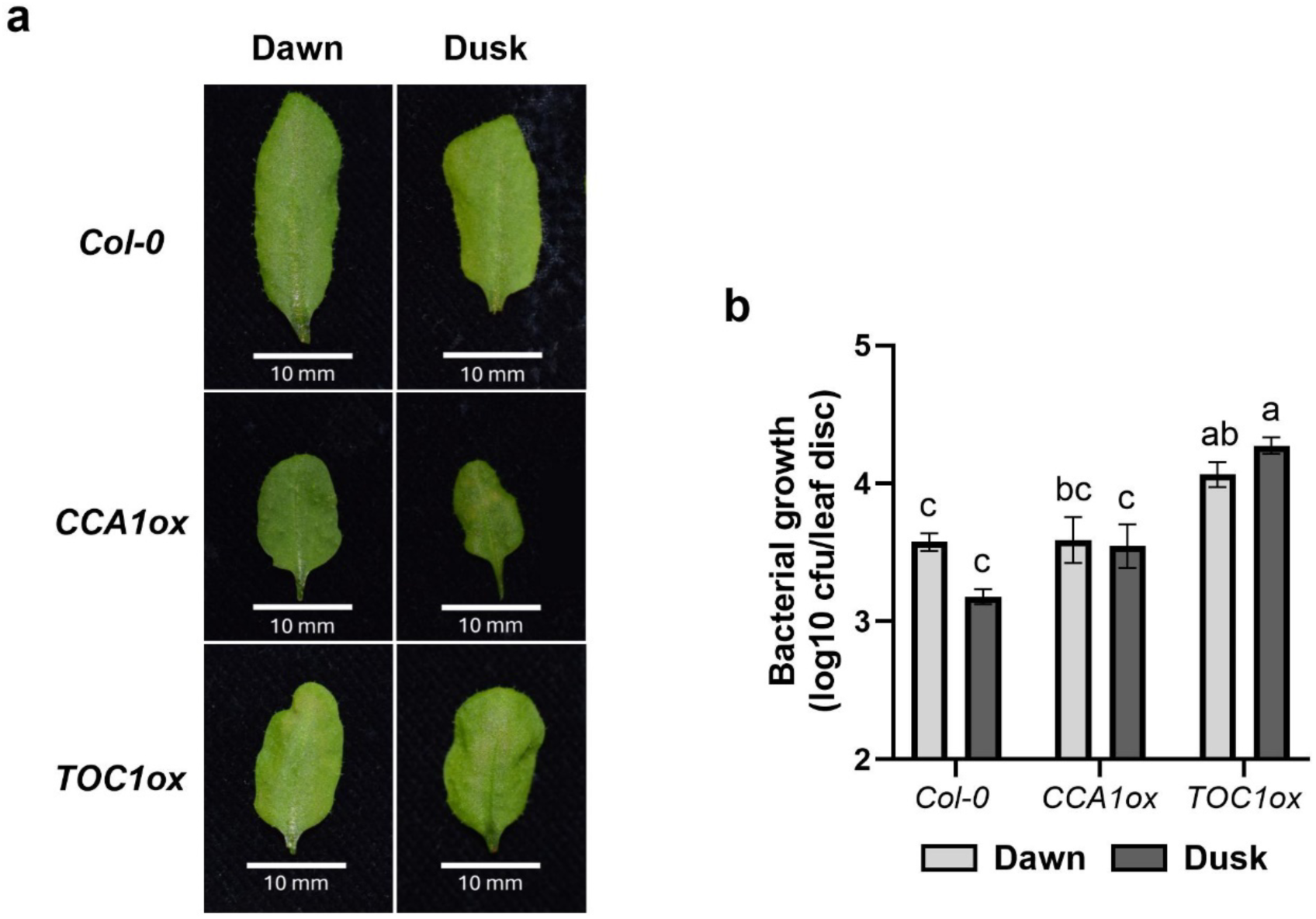
*TOC1ox* is more susceptible to *Pst* DC3000 *hrcC* infection. *Col-0*, *CCA1ox* and *TOC1ox* plants were grown under 12/12 light/dark conditions for 4 weeks. At dawn on day 28 (LL0) plants were released into constant light conditions. At LL24 or LL36 the plants were infected with *Pseudomonas siringae pv. tomato (Pst)* DC3000 *hrcC*. The level of bacterial growth was measured 5 days post infection. (a) Leaves represent the visual median level of infection 5 days post infection for each genotype and time. (b) The infection level 5 days post infection expressed by colony forming units (CFU; y-axis is logarithmic) in leaf disk extracts. Error bars represent mean ± SEM (n = 8 individual leaves). A two-way ANOVA and Tukey’s HSD Test were preformed to analyse the differences in bacterial growth between time points and genotypes.

Overall, these results suggest that regulated *TOC1* levels and not general clock rhythmicity are integral for PTI immunity with or without the interference of effectors.

The letters indicate significant difference between groups (p < 0.05). The data shown here are from a single experiment representative of 3 independent experiments.

### Flg22 or elf18 treatment shortens the period of *CCA1* and *TOC1* rhythms

Our investigation to this point has focused on circadian clock regulation on the immune system, however evidence of the reciprocal influence also exists. Previous research found that flg22 treatment shortened the period of *CCA1* rhythms ^30^. We wanted to investigate whether the period shortening effect was dose dependent and whether elicitor treatment also affected *TOC1* rhythms. We also investigated the effect of bacterial elicitor elf18. Leaf disks of plants expressing firefly luciferase (LUC) driven by the rhythmic promoters of either *CCA1* or *TOC1* (*CCA1pro:LUC* and *TOC1pro:LUC*, respectively) were subjected to a range of flg22 or elf18 concentrations.

Flg22 treatment induced dose-dependent period shortening of *CCA1* (Figure 8a-d) and *TOC1* (Figure 8i-l) rhythms under constant light conditions. Elf18 treatment also induced period shortening in both *CCA1* (Figure 8e-h) and *TOC1* (Figure 8m-p), though it was not dose dependent. These findings indicate that the detection of PAMPs modulates circadian rhythms, potentially to facilitate an effective defence response.

**Figure 8.**
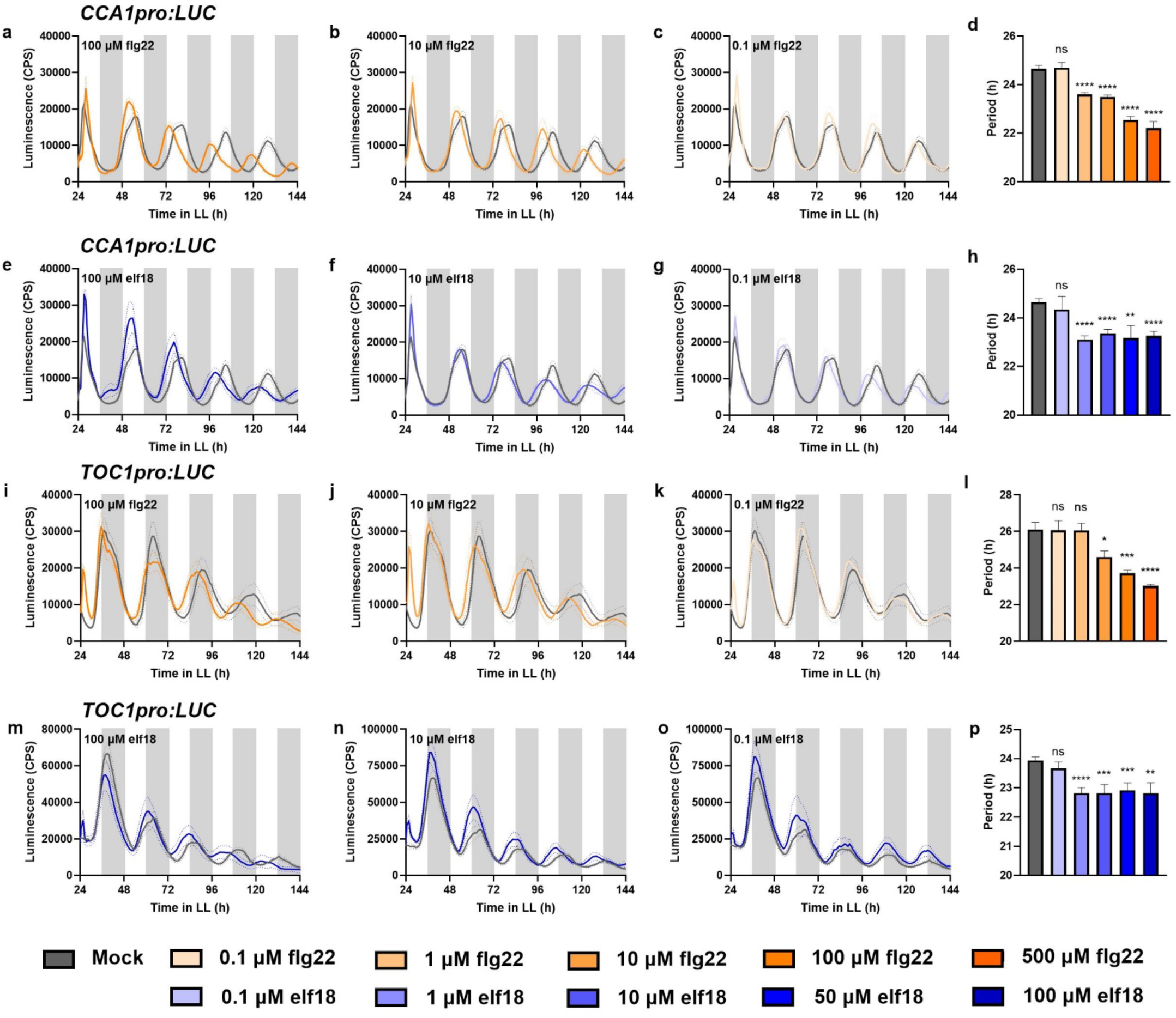
Flg22 or elf18 treatment shortens the period of *CCA1* and *TOC1* rhythms. Promoter activity of *CCA1* and *TOC1* was observed by measuring luminescence in *CCA1pro:LUC* and *TOC1pro:LUC* leaf disks, respectively. Leaf disks were treated with mock (ddH_2_O) or a range of concentrations of either flg22 (500 μM, (a,i) 100 μM, (b,j) 10 μM, 1 μM and (c,k) 0.1 μM) or elf18 ((e,m) 100 μM, 50 μM, (f,n) 10 μM, 1 μM and (g,o) 0.1 μM). Only luminescence traces for 100, 10 and 0.1 μM of each elicitor are shown here to represent the overall results. The mean period for all elicitor concentrations (d, h, l, p) was calculated from the 24 - 120 h time window. Unpaired t-tests were performed between each treated data set and the mock data set: *p<0.05, **p<0.01, ***p<0.001, ****p<0.0001. Error bars indicate mean ± SEM (n = 8). Data are from single experiments, representative of 3 independent repeats each.

### Flg22- and elf18-induced period shortening depends on their cognate receptors

PAMPs are detected by specific pathogen recognition receptors at the plasma membrane. Therefore, we next investigated whether the presence of the pathogen recognition receptor associated with recognising flg22 and elf18 (FLS2 and EFR, respectively) were required for this period shortening effect on the clock. *CCA1pro:LUC* and *TOC1pro:LUC* plants were crossed with *fls2* and *efr1* mutant plants separately creating 3 new lines: *CCA1pro:LUC* in the *fls2* or *efr1* background and *TOC1pro:LUC* in the *efr1* background. We were unable to produce homozygous crosses between *TOC1pro:LUC* and *fls2* plants so this line was not analysed in this study. We found that the period shortening induced by flg22 was not observed in *CCA1* rhythms in the *fls2* background (Figure 9a-d). Additionally, elf18 induced period shortening was not observed in *CCA1* (Figure 9e-h) or *TOC1* (Figure 9i-l) rhythms in the *efr1* background. We conclude that the pathogen recognition receptor responsible for detecting of the elicitor is required for the subsequent effect on the clock, and therefore that the effects observed in Figure 8 are specific and mediated by the known pathways that mediate PAMP perception.

**Figure 9.**
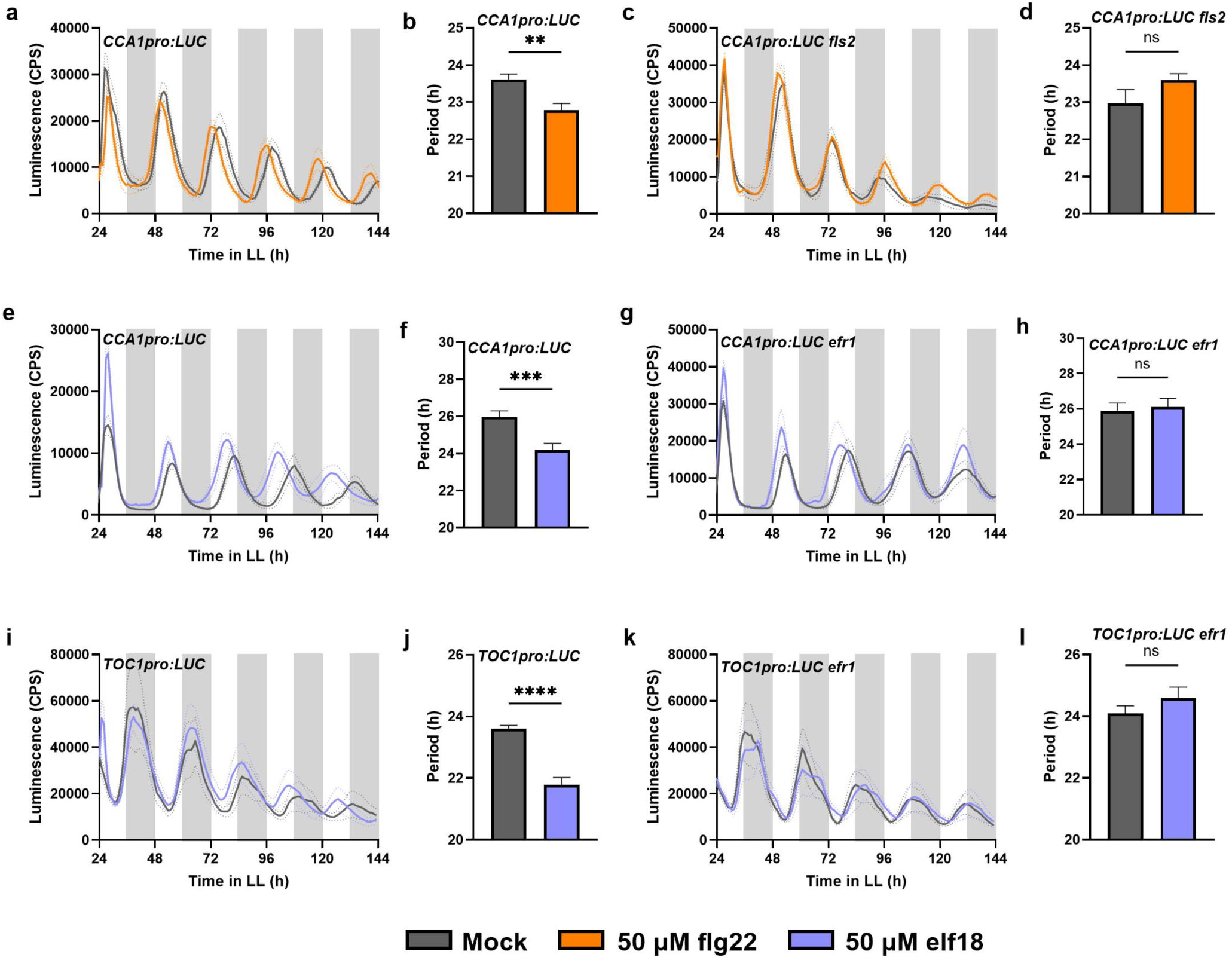
The period shortening eAect of flg22 or elf18 is lost in the *fls2* or *efr1* background, respectively. Promoter activity of *CCA1* and *TOC1* was observed by measuring luminescence in *CCA1pro:LUC* and *TOC1pro:LUC* leaf disks, respectively, in the wild type (a,e,i) or *fls2* (c) or *efr1* (g,k) background. Leaf disks were treated with mock (ddH_2_O) or 50 μM flg22/elf18. The mean period of the wild type (b,f,j), *fls2* (d) and *efr1* (h,l) leaf disks was calculated from the 24 - 120 h time window. Unpaired t-tests were performed between the treated data set and the mock data set: *p<0.05, **p<0.01, ***p<0.001, ****p<0.0001. Error bars indicate mean ± SEM (n = 8). Data are from single experiments, representative of 3 independent repeats each.

## Discussion

Plant pathogens are responsible for huge losses in annual crop yield worldwide and plant disease outbreaks are increasing in frequency, threatening food security ^47–49^. Many challenges facing crops are governed by the light dark cycle of our planet, including pathogen virulence and activity ^5,50–52^. A functioning circadian clock improves plants resistance to pest and pathogens both before and after harvest, suggesting that further understanding of this relationship could yield useful results for improving crop health ^50,53^. In this study, we examined the relationship between PTI signalling and the circadian clock in *Arabidopsis,* through investigation of PAMP- triggered defence pathways.

The results we have presented here indicate that the transcript levels of early immune gene *FRK1* is visually rhythmic under basal conditions and significantly rhythmic when induced by bacterial elicitor flg22, but not elf18. We found that basal *FRK1* transcript levels peaked during the subjective day whereas flg22-induced *FRK1* transcript levels peaked during the subjective night/around subjective dusk. Previous studies reported circadian rhythmicity in various genes involved in the flg22 recognition pathway under basal conditions including *FLS2* and some *MAPKs* ^27^. Similarly to our results, Bhardwaj et al. ^27^ found that basal expression of flg22 pathway genes peaked during the subjective day. This coincides with when stomata are open, which pathogens can use as an entry point making infection more likely ^39^. The shift in the peak transcript levels of *FRK1* that we observed after flg22 induction indicates that the circadian clock gates this response, responding more severely during the subjective night/around subjective dusk when pathogen infection may be less anticipated as stomata are closed. Furthermore, we show that the temporal difference in flg22-induced transcript levels of *FRK1* are disrupted in *CCA1ox* plants and *cca1lhy* mutants. This suggests that the rhythmicity of the clock and regulated expression *CCA1* are important to maintain the temporal difference in levels of flg22-induced *FRK1* transcript levels between subjective dawn and dusk. Significantly, we show that an increase in *FRK1* transcript levels in response to flg22 was not observed in *TOC1ox* plants at either subjective dawn or dusk. This suggests that the overexpression of *TOC1* suppresses this defence response to flg22 detection. In a similar experiment, we found that the transcript levels of flg22 pathogen recognition receptor *FLS2* were significantly elevated following flg22 treatment in *Col-0* and *CCA1ox* plants but not in *TOC1ox* plants. *toc1-101* plants did not exhibit consistently higher flg22-induced *FRK1* transcript levels at subjective dusk. This disruption, like the disruption observed in *cca1lhy* mutants, could be due to the short period exhibited by these mutants. Furthermore, the *FRK1* transcript levels after flg22 treatment in *toc1-101* plants were not hugely different to *Col-0.* Additionally, ChIP-Seq data indicates that TOC1 does not directly bind *FLS2* ^8^. Together, this suggests that *TOC1* is not directly responsible for the suppression of *FRK1* and *FLS2* in response to flg22 treatment, but rather its overexpression has regulatory effects on some other component that controls the expression of these PTI genes. Additionally, levels of flg22-induced *FLS2* transcript in *CCA1* were consistently higher than those observed in *Col-0.* This could suggest that the overexpression of *CCA1* promotes flg22-induced responses, however this was not a result we observed in the rest of our data.

In addition to circadian regulation of early defence genes, we demonstrate circadian regulation of the ROS burst in response to flg22 treatment. Wild type (*Col-0*) plants produced a larger ROS burst when challenged with flg22 at subjective dusk compared to subjective dawn. This distinct difference in response was observed in *cca1lhy* mutants but not in *CCA1ox* plants. Interestingly, we did not observe any flg22-induced ROS burst in *TOC1ox* plants at either dawn or dusk. These results again indicate that the flg22-induced early immune response is stronger at subjective dusk than subjective dawn. It appears that only arrhythmicity of the clock through overexpression of *CCA1* and not loss of *CCA1* disrupts the difference in the response of the ROS burst between subjective dawn and dusk. Lai et al.^29^ found that *CCA1ox* plants had lower basal ROS levels and attenuated induction of ROS genes in response to oxidative stress. This suppression of overall ROS activity by the overexpression of *CCA1* could be causing the loss of the larger ROS burst at dusk that we observe in wild type (*Col-0*). The complete lack of a ROS burst response to flg22 in *TOC1ox* plants suggests that overexpression of *TOC1* also supresses this early defence response. As *toc1-101* does not appear to differ from wild type in phenotype, this suggests again that it is not direct suppression by *TOC1* but indirect suppression through another element of the clock that *TOC1* negatively regulates. Korneli et al. ^28^ also found circadian control of the flg22 induced ROS burst, however they reported that ROS production was higher in plants challenged in the subjective morning versus subjective evening. This result differs to ours, possibly due to significant differences in methodology: they cut leaf disks only 6 hours before treatment. Research has shown that damage associated molecular patterns (DAMPs) may induce increased levels of ROS production for up to 6 hours after damage ^54–56^. To avoid this problem, we cut leaf disks 24 hours before treatment. Korneli et al. ^28^ furthermore found that arrhythmicity of the clock abolished the distinct difference in levels of ROS production between morning and evening through analysis of *lux* plants which display reduced rhythmicity of clock genes ^57^. *LUX* is a member of the evening complex (EC), which is negatively regulated by *TOC1* ^7,11,13^. Therefore, it may be worth investigating other elements of the EC as candidates for regulation of the flg22-induced ROS burst and other flg22-induced PTI responses, as they are regulated by *TOC1.* Alongside Lai et al. ^29^, other research has also shown circadian clock involvement in oxidative stress tolerance and ROS production ^58,59^, suggesting an extensive involvement of the clock in ROS homeostasis. Overall, our results continue to indicate that the clock enhances elements of the PTI response to flg22 at subjective dusk/night and that the overexpression of *TOC1* has hugely detrimental effects on flg22-induced early defence responses.

PTI pathways involve rapid responses, some of which we have investigated in this study; the peak of the ROS burst occurs within 10 minutes ^28^ and increased transcript levels of early immune genes can be detected within 1 hour ^35^. These are specific responses to PAMP detection that function together to defend the plant against infection. To determine the cumulative effects of clock interference with these processes, it is useful to observe later immune responses such as susceptibility to infection. However, the later actions of PTI are more difficult to investigate, as the pathogen attempts to facilitate infection by injecting effectors which affect host susceptibility ^40,41^. By infecting plants with the *Pst* DC3000 *hrcC* mutant and *Pst* DC3000 wild type we are able to observe how the clock interacts with PTI during virulent infection with and without manipulation by effectors ^45,46^. We observed no difference in infection levels between subjective dawn and dusk for *Col-0, CCA1ox* or *TOC1ox* plants infected with *Pst* DC3000 *hrcC*, suggesting that there is not clock gating of PAMP triggered resistance at these times. *Col-0* showed increased susceptibility to infection with *Pst* DC3000 at subjective dusk compared to dawn. *TOC1ox* plants did not shown any difference in infection level between subjective dawn and dusk and were significantly more susceptible to both *Pst* DC3000 *hrcC* and *Pst* DC3000 compared to *Col-0*. *CCA1ox* plants displayed inconsistent temporal differences in susceptibility to *Pst* DC3000. Bhardwaj et al. ^27^ similarly found that *Col- 0* was most resistant to infection with *Pst* DC3000 in the subjective morning and most susceptible during the subjective night. In *CCA1ox* plants and *elf3-1* mutants, both of which are arrhythmic ^60–62^, they found that there was no difference in susceptibility between subjective morning and night. This suggests that clock arrhythmicity disrupts the increased resistance observed in the subjective morning. Our observations of *CCA1ox* plants were inconsistent, with two repeats displaying temporal differences in susceptibility similar to *Col-0* and a final experiment showing no temporal difference. There are some differences in experimental set up that could have contributed to this discrepancy: Bhardwaj et al. ^27^ grew plants under 16/8 light/dark conditions and thus had different sample times. Neither our study nor theirs investigates infection levels in *CCA1ox* at multiple regular time points over 24-48 hours. We suggest that levels of *CCA1* are not consistent or regulated in the *CCA1ox* plants and therefore create an inconsistent result between experiments. Therefore, we cannot conclude that resistance to *Pst* DC3000 in *CCA1ox* is rhythmic or not rhythmic, only that it displays inconsistent temporal differences in susceptibility in our study and does not display these differences in the study by Bhardwaj et al. ^27^. On the other hand, our results consistently show the clock does not enhance overall resistance to *Pst* DC3000 at subjective dusk, while we do observe increased flg22-induced *FRK1* transcript levels and a stronger flg22-induced ROS burst at that time. Rather, *Col-0* plants appear to be more susceptible to infection at subjective dusk. In a previous study, we found that *CCA1* expression was essential for SA-induced resistance to *Pseudomonas syringae* pv. *manculicola,* suggesting that *CCA1* may be more significantly involved in later SA-induced immunity than PTI ^63^. Significantly, *TOC1* overexpression has negative consequences for PTI, with *TOC1ox* plants displaying increased susceptibility to *Pst* DC3000 at subjective dawn and increased susceptibility to the *hrcC* mutant at both subjective dawn and dusk. It appears that the overexpression of *TOC1* must disrupt the regulation of multiple components that are essential for PTI. It has been shown that *CCA1* confers heterosis for bacterial defence in *Arabidopsis* hybrids (*Col-0* x *Sei-0*) without costs to growth vigour. This suggests that *CCA1* is potentially a positive regulator of immunity, however, it is likely not that simple as this study and our previous study show that immunity to bacterial infection is not improved in plants overexpressing *CCA1* or worsened in plants missing *CCA1* (*cca1lhy*) ^63^. Similarly, we are not suggesting that *TOC1* is a direct negative regulator of immunity, but that it must be involved in the regulation of an intermediary component and, only when overexpressed, has hugely detrimental effects on the modulation of PTI immune responses.

In the final section of this study, we investigated the effect of PTI signalling on clock gene rhythms. We found that both flg22 and elf18 induced period shortening of both *CCA1* and *TOC1* rhythms, complementing previous research by ^30^ that showed flg22-induced period shortening of *CCA1* rhythms. In addition to this, we found that the flg22-induced period shortening was dose dependent, but elf18-induced period shortening was not. These results, combined with the rhythmicity in flg22-induced *FRK1* transcript and lack of rhythmicity in elf18-induced *FRK1* transcript levels, suggest that there may be a more significant link between the circadian clock and the flg22 recognition pathway compared to the elf18 recognition pathway. We also showed that the elicitor induced period shortening was dependent on the expression of the corresponding pathogen recognition receptor (*FLS2* for flg22 and *EFR* for el18), indicating that without detection at the membrane, these elicitors do not affect the clock. Together, these results demonstrate that plants might use perception of flg22 levels to modulate the clock according to the magnitude of the detected threat. Similar research investigating the effect of immune hormone salicylic acid (SA) on the clock has also shown period shortening of clock gene rhythms following SA treatment ^63,64^. This indicates that various immune signalling pathways may use the circadian clock in a similar way to regulate other cellular processes in order to promote an efficient defence response.

Overall, the work in this study indicates that the circadian clock gates individual flg22-induced early immune responses, resulting in a more severe response to threat in the subjective evening and night versus subjective morning. However, overall susceptibility to infection is increased at subjective dusk compared to dawn. Consistent through all our results, *TOC1* overexpression has a profound effect on PTI processes. In all cases in this study, overexpression of *TOC1* has resulted in suppression of the PAMP-triggered defence responses observed in wild type plants and increased susceptibility to infection. As no significant response was observed in *toc1-101* mutants, these results indicate that the overexpression of *TOC1* disrupts the activity of other factors controlling immunity that remain rhythmic in the *toc1-101* mutant. To determine the intermediate components, it could be useful to investigate the involvement of clock genes regulated by *TOC1* such as members of the evening complex. We also demonstrate reciprocal regulation of the circadian clock by the PAMP-triggered immune signalling. The findings presented in this study greatly further our understanding of the interplay between the circadian clock and early immune responses. Continued study of how *TOC1* overexpression supresses PAMP-triggered immune responses could aid in uncovering the mechanisms underpinning the regulation of PTI in plants.

## Methods

### Plant materials and growth conditions

*Arabidopsis thaliana (Arabidopsis)* seeds were sterilised for 5 minutes in 70% ethanol followed by 5 minutes in 20% bleach (0.25% sodium hypochlorite, Cleanline) and washed 3 times in ddH2O. Seeds were then stratified in ddH2O at 4 ᵒC for 3 days before sowing.

*Arabidopsis* plants were grown on autoclaved soil for 4 weeks, unless otherwise stated, in growth cabinets (Percival Scientific Inc.) set at constant 21ᵒC with a light intensity of 70-100 µmoles m^-2^ s^-1^ (Fusion 18W T8 2ft Triphosphor Fluorescent Tube 4000K) under 12 h light/12 h dark. All wild type and mutant plants were from the *Col-0* genetic background and were described previously: *fls2* ^65^*; efr1* ^34,66^*; CCA1ox* ^60^; *TOC1ox* ^67^; *cca1lhy* ^30^; *toc1-101* ^68^; *CCA1pro:LUC* ^69^ and *TOC1pro:LUC* ^70^.

*CCA1pro:LUC fls2, CCA1pro:LUC efr1* and *TOC1pro:LUC efr1* plants were made by crossing *CCA1pro:LUC* and *TOC1pro:LUC* plants with *fls2* or *efr1* mutant plants. To obtain homozygous lines of *fls2* (SAIL_691C04) or *efr1* (SALK_044334), plants of the F1 and F2 generations were evaluated by polymerase chain reaction (PCR) genotyping with the corresponding left border primer to detect the T-DNA insertion (LB2 for SAIL and LBb1.3 for SALK) and the gene specific right primer (SIGnal T-DNA Express, http://signal.salk.edu/tdnaprimers.2.html). PCR products were visualized on 1% agarose gel.

Homozygous *fls2* or *efr1* lines were tested for homozygosity of the luciferase transgene (either *CCA1pro:LUC* or *TOC1pro:LUC*) in F3. A minimum of 50 seeds of each line were visualized for luminescence by sowing one seed per well, on top of 200 μl 0.5 MS in black 96-well plates (Greiner bio-one, 655075). Seedlings were grown for 10 days in an incubator (Sanyo) under long-day conditions. 20 μl of 1 mM D-luciferin (Biosynth AG) was added to each well and luminescence was visualized using the ALLIGATOR luminescence imaging system by exposing the camera for 8 minutes. Ratios of luminescence were assessed: homozygous lines which displayed luminescence in all seedlings were used for subsequent experiments.

### Elicitor treatment for quantitative PCR in wild type over 48 hours

Wild type plants (*Col-0*) were grown on MS agar plates under 12 h light/12 h dark conditions at 21ᵒC with a light intensity of 70-100 µmoles m^-2^ s^-1^ (Fusion 18W T8 2ft Triphosphor Fluorescent Tube 4000K). After 12 days, the seedlings were released into constant light (LL). Plates were removed from the growth cabinet and sprayed with 1 µM flg22, 1 µM elf18 or mock (ddH_2_O) at 2- hour intervals beginning at LL24 and finishing at LL72. After spraying, the plates were put into separate growth cabinets from the untreated plates. The seedlings were harvested and frozen in liquid nitrogen 2 hours after treatment.

### Elicitor treatment for quantitative PCR in clock mutants

Wild type (*Col-0*), *CCA1ox, cca1lhy* and *TOC1ox* plants were grown on MS agar plates under 12 h light/12 h dark conditions at 21ᵒC with a light intensity of 70-100 µmoles m^-2^ s^-1^ (Fusion 18W T8 2ft Triphosphor Fluorescent Tube 4000K). Plates were placed under split entrainment conditions in separate growth cabinets, aligning the time of dawn and dusk between the two cabinets. After 12 days, the seedlings were released into LL. Plates were sprayed with 1 µM flg22, 1 µM elf18 or mock (ddH2O) at the same time, coinciding with either subjective dawn (LL24) or dusk (LL36).

The seedlings were harvested and frozen in liquid nitrogen 2 hours after treatment.

### RNA extraction and cDNA synthesis

Seedlings were manually ground to a fine powder in liquid nitrogen, before being homogenized in RNA extraction buffer (100 mM LiCl, 100 mM Tris pH 8, 10 mM EDTA, 1% SDS). An equal volume of phenol/chloroform/isoamyl alcohol (25:24:1) was added, followed by votexing and centrifuging at 13,000 rpm for 5 min. The aqueous phase was transferred to a tube containing an equal volume of chloroform/isoamyl alcohol (24:1), followed by vortexing and centrifuging at 13,000 rpm for 5 min. This step was repeated before the aqueous phase was transferred to a tube containing 1/3 volume of 8 M LiCl and incubated overnight at 4ᵒC. The samples were centrifuged at 13,000 rpm at 4ᵒC for 15 min. The supernatant was removed and the pellet was washed twice in ice-cold (-20ᵒC) 70% ethanol. The pellet was then dissolved in 400 µl H2O for 30 min on ice. 40 µl of NaAc (pH 5.3) and 1ml of ice-cold 98% ethanol was added and the solution was incubated for at least 1 h at -20ᵒC. The tubes were centrifuged at 13,000 rpm for 15 min at 4ᵒC and the pellet was washed twice with ice-cold 70% ethanol. The final pellet was dissolved in 50 µl H_2_O. The resulting RNA was quantified using a NanoDrop spectrophotometer and diluted to standardize the concentrations across all samples. SuperScript II reverse transcriptase (Invitrogen) was used to perform reverse transcription according to the manufacturer’s instructions.

### Quantitative PCR (qPCR)

cDNA was diluted 20-fold and qPCR was performed in a 5 µl reaction using SYBR Green and gene specific primers on a QuantStudio 5 PCR machine. All qPCR experiments used housekeeping gene *SFP* (*SAND Family Protein)* to normalize the data ^71^.

### Circadian rhythmicity analysis of transcript levels

Elicitor-induced *FRK1* transcript levels in wild type over 48 hours (Supplementary Figure 1) were assessed for circadian rhythmicity using the JTK_cycle algorithm with empirical calculation of *p*- values (eJTK) in BioDare2^36,72^. The mean transcript levels of four technical replicates were used for analysis with parameters set as eJTK Classic, linear detrending of the input data and p<0.05.

### Measurement of the apoplastic ROS burst

*Col-0, CCA1ox, cca1lhy* and *TOC1ox* were grown on autoclaved soil under split entrainment conditions as described previously in: Elicitor treatment for quantitative PCR in clock mutants. At 4 weeks old, 4 mm leaf disks were cut from leaves 5-6 and placed adaxial side up into the wells of a black, flat bottomed 96-well plate (Greiner bio-one, 655076). The wells contained 200 µl of ddH_2_O ^73^. The plate was transferred to constant light conditions (70-100 µmoles m^-2^ s^-1^, Fusion 18W T8 2ft Triphosphor Fluorescent Tube 4000K). At dawn the following day (coinciding with LL24 and LL36) the water was carefully removed from the wells and replaced with 100 µl of elicitation solution (0.2 µM luminol [dissolved in 200 mM KOH], 20 µg/ml horseradish peroxidase and 0.1 µM flg22 for treated wells). The plate was sealed with a clear plastic lid (Revvity Health Sciences Inc, 6050185) and luminescence was measured immediately by a LB942 Tristar2 plate reader (Berthold Technologies Ltd) for 1 second per well for 60 minutes. Results were analyzed using GraphPad Prism.

### *Pst* DC3000 and *Pst* DC3000 *hrcC* disease assay

*Col-0, CCA1ox* and *TOC1ox* plants were grown on soil under 12/12 light/dark conditions for 4 weeks. Plants were grown under split entrainment conditions in separate growth cabinets, aligning the time of dawn and dusk between the two cabinets. On day 28, plants were transferred to constant light conditions (LL). At dawn the following day, subjective dawn or subjective dusk depending on the cabinet, one leaf per plant was pressure infiltrated with *Pseudomonas* using a 1 ml syringe at OD_600_= 0.01. The bacteria were left to infect the plants for 3 days for *Pst* DC3000 or 5 days for *Pst* DC3000 *hrcC*. Leaf disks were then harvested from the infiltrated leaves and homogenized with 10 mM MgCl_2_. The leaf disk extract was diluted ten-fold 5 times and these dilutions were plated onto LB agar plates (10 mM MgCl_2_ and 50 µg/ml rifampicin). Plates were incubated for 2 days at 28ᵒC before calculating colony forming units (CFU) per leaf disk.

### Bioluminescence assay of leaf discs

*CCA1pro:LUC, TOC1pro:LUC, CCA1pro:LUC fls2, CCA1pro:LUC efr1* and *TOC1pro:LUC efr1* were grown on soil under LD conditions (see plant materials and methods). At 4 weeks old, leaf disks were taken from leaves 5-6 and placed in the wells of a white, flat-bottom 96-well plate (Greiner bio-one, 655075) with the adaxial side up. Each well contained 180 µl of filter sterilized imaging solution: 0.5 MS pH 7.5, 50 µg/ml ampicillin, 1.5 mM D-luciferin and ddH_2_O to the desired final concentration. The plate was sealed with a clear, gas permeable lid (4titude, 4ti- 0516/96). Luminescence was measured by a LB942 Tristar2 plate reader (Berthold Technologies Ltd) every 50 minutes for 3 seconds per well. Leaf disks were kept under continuous red (630 nm) and blue (470 nm) LED light at 17.5 µmoles m^-2^ s^-1^ each at 20-21ᵒC. After 24 h in constant light (LL) plants were treated with 20 µl mock (ddH_2_O) or an elicitor solution to the desired concentration. The plate is sealed with a new clear, gas permeable lid and returned to the plate reader. Results were analysed using GraphPad Prism and Biodare2^72^, and period was estimated using the Fast Fourier Transform-Non-Linear Least Squares (FFT-NLLS) function.

## Data availability Statement

The Supplementary Figure 1 is available within the paper and its Supplementary Information file. The datasets used and/or analysed during the current study are available from the corresponding author on reasonable request.

## Acknowledgements

We would like to thank the members of the van Ooijen Lab and the Spoel Lab at the University of Edinburgh for their assistance with this study. We would like to thank Karen Halliday for kindly giving us the clock mutant lines and luciferase lines used in this study. We would like to thank the lab of Cyril Zipfel for the donation of the *fls2* and *efr1* lines used in this study. Fanyu Zheng assisted in generating the samples for Supplemental Figure 1, and Samantha Cargill contributed to generating plant material.

This research was carried out with resources provided by the Edinburgh Plant Growth Facility, a specialist service provider for plant growth at the University of Edinburgh.

OJPF was support for the research of this work from the Biotechnology and Biological Sciences Research Council EASTBIO Doctoral Training Program (BBSRC, BB/M010996/1).

## Author contributions

GvO, OJPF, and SHS conceptualised and designed the research. OJPF performed all experiments and data analysis and prepared the figures. OJPF and GvO wrote the manuscript. All authors read and approved the final manuscript.

