## Supplemental Figure 1 for "TOC1 supresses PAMP-triggered immunity in *Arabidopsis*"

### Supplementary Figures

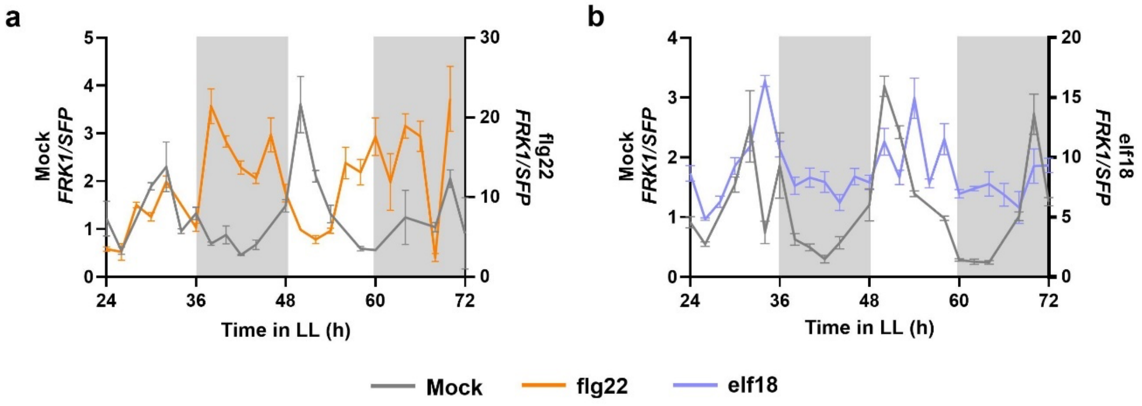


**Supplementary Figure 1. Flg22- but not elf18-induced *FRK1* transcript levels are rhythmic.** *Col-0* seedlings were grown on MS plates under 12/12 light/dark conditions for 12 days before being released into constant light (LL). Every 2 hours, plates were sprayed with 1 µM flg22 (a, orange), elf18 (b, blue) $ddH_2O$ (mock, grey). Transcript levels of *FRK1* was measured 2 hours post treatment. The data points are the mean ± SEM of 4 technical replicates. The data shown here are from a single experiment of which there are no repeats.
